# Metabolic network construction reveals probiotic-specific alterations in the metabolic activity of a synthetic small intestinal community

**DOI:** 10.1101/2023.03.29.534679

**Authors:** Jack Jansma, Anastasia Chrysovalantou Chatziioannou, Kitty Castricum, Saskia van Hemert, Sahar El Aidy

## Abstract

The gut microbiota plays a crucial role in maintaining overall health and probiotics have emerged as a promising microbiota-targeted therapy for improving human health. However, the molecular mechanisms of probiotics action in general and the targeting of small intestinal microbiota by probiotics are not well understood. To address this, we constructed a synthetic community of three species, which resembles the upper small intestinal microbiota. Our results indicate that probiotic supplementation can directly affect the metabolism of the community, resulting in colonization resistance in a probiotic specific manner. Supplementation with *Streptococcus thermophilus* led to increased lactate production and a decrease in pH, while *Lactobacillus casei* supplementation increased the resistance to perturbations and nutrient utilization without affecting lactate production or pH. Additionally, when combined with kynurenine, *Lactobacillus casei* enhanced the kynurenine pathway metabolism resulting in elevated kynurenic acid levels and possibly indirect colonization resistance. Overall, our study reveals how selecting probiotics with distinct functional capacities can unlock the full potential of microbiota-targeted therapies.

**Importance:** The development of probiotic therapies targeted at the small intestinal microbiota represents a significant advancement in the field of probiotic interventions. This region poses unique opportunities due to its low number of gut microbiota, along with the presence of heightened immune and metabolic host responses. However, progress in this area has been hindered by a lack of detailed understanding regarding the molecular mechanisms through which probiotics exert their effects in the small intestine. Our study, utilizing a synthetic community of three small intestinal bacterial strains and the addition of two different probiotic species, and kynurenine as a representative dietary or endogenously produced compound, highlights the importance of selecting probiotic species with diverse genetic capabilities that complement the functional capacity of the resident microbiota, or alternatively, constructing a multispecies formula. This approach holds great promise for the development of effective probiotic therapies and underscores the need to consider the functional capacity of probiotic species when designing interventions.

## Introduction

In recent decades, there has been a growing interest in investigating the gut-microbiota and its significant roles in maintaining human health [1, 2]. The gut-microbiota is composed primarily of *Bacteroidaceae*, *Prevotellaceae*, *Rikenellaceae*, *Lachnospiraceae* and *Ruminococcaceae*, which reside mainly in the colon [3]. In contrast to the colon, the small intestinal microbiota is highly dynamic and location-specific. For example, *Streptococcaceae* and *Veilonellaceae* are abundant in the duodenum [4], while *Streptococcaceae*, *Carnobacteriaceae* and *Actinomycetaceae* prevail in the jejunum [5] and *Lactobacillaceae*, *Erysipelottrichaceae*, and *Enterobacteriaceae* are dominant in the ileum [6]. The small intestinal microbiota plays a crucial role in regulating the metabolic, endocrine, and immune functions of their hosts [7–9]. Consequently, altering the equilibrium between the host and the microbiota can negatively affect the health of the host [10, 11]. Thus, to re-establish this equilibrium and enhance the health of the host, multiple microbiota-targeted therapies, such as probiotics, are employed [12].

Probiotics are defined as live microorganisms that, when administered in adequate amounts, confer a health benefit on the host [13]. The development of probiotic therapies aiming at targeting the small intestinal microbiota poses a significant challenge due to the limited access to the human small intestine. Currently, investigations into the effect of probiotics on the small intestine are carried out either directly on animal models [9, 14] or indirectly through small clinical trials that lack detailed molecular mechanisms [15]. This lack of understanding hampers the development of novel probiotic therapies. In recent years, it has been demonstrated that the microbiota retrieved from stoma effluent exhibits similar compositional and behavioral characteristics with the healthy ileal microbiota [16]. These findings suggest that the stoma effluent-derived microbiota may serve as a promising source for probiotic research [17, 18]. However, the utilization of stoma effluent is limited for the study of the upper small intestine. To address this limitation, there is a need for the construction of synthetic bacterial communities to bridge this research gap [19]. Synthetic bacterial communities are bacterial mixtures produced through co-culturing of selected bacteria in controlled environments [20]. These communities can comprise a range from two [21] to over one hundred species [22]. Synthetic communities can be subjected to perturbation through manipulation of growth parameters or the introduction of new bacterial species, and can also be supplemented with specific compounds such as prebiotics [23]. Analysis of the community’s molecular response allows a deeper understanding of the underlying molecular mechanisms [24], optimization of metabolites’ production [25] and assessment of emergent properties such as stability or resistance to perturbations [26, 27]. This methodology has the potential to guide the development of novel microbiota-targeted therapies.

In humans, the kynurenine pathway is the primary route for the metabolism of free tryptophan, and it plays a critical role in the generation of energy in the form of nicotinamide adenine dinucleotide (NAD^+^), immune system regulation [28] and the production of the neuroactive compounds quinolinic and kynurenic acid (KYNA) [29]. Increased tryptophan metabolism via the kynurenine pathway in the gut is associated with obesity and alterations in the microbiota [29]. G-protein coupled receptor 35 (GPR35), which is activated by KYNA, is abundantly expressed in the intestine, where it plays a role in immune regulation and epithelial barrier function [30]. The concentration of KYNA in the gut lumen increases from the proximal to the distal part of the intestine, indicating production by the gut microbiota [31, 32]. Several gut bacteria can produce compounds from the kynurenine pathway [32, 33] and probiotics supplementation have been shown to reduce intestinal inflammation and improve intestinal barrier function accompanied by reduced kynurenine but increased KYNA levels [34, 35]. However, the precise effects of kynurenine pathway metabolism within the microbiota remain unknown.

In the present study, a dynamic correlation-based network approach was utilized in conjunction with multivariate analysis of experimental data obtained from chip-based digital polymerase chain reaction (c-dPCR), proton-nuclear magnetic resonance (^1^H-NMR) and high-performance liquid chromatography-ultraviolet (HPLC-UV). The aim was to examine the impact of introducing probiotic species on the general metabolic activity and the kynurenine pathway activity of a simplified community consisting of three intestinal isolates, which represent the small intestinal microbiota.

## Material and methods

### Bacteria and growth media

Bacteria were selected based on their abundance in the upper small intestine, ability to metabolically affect the kynurenine pathway or probiotic activity in the small intestine. *Escherichia coli* DSM11250 was obtained from the German collection of microorganisms and cell cultures; *Pseudomonas fluorescens* MFY63 was obtained from the laboratory of microbiology signals and microenvironment [36]; *Streptococcus salivarius* HSISS4 was obtained from the laboratory of host-microbe interactomics [37]; *Streptococcus thermophilus* W69 and *Lactobacillus casei* W56 were obtained from Winclove probiotics. All individual bacterial and community growth experiments were performed using enriched beef broth (EBB) [38].

### Bacterial kynurenine pathway metabolism

To investigate kynurenine pathway metabolism by the bacterial strains, 3 mL EBB was inoculated from a glycerol stock stored at –80 °C and grown overnight at 37 °C. *E. coli* and *P. fluorescens* were grown shaking at 220 RPM, *S. salivarius*, *S. thermophilus* and *L. casei* were grown without shaking. Overnight grown pre-culture were diluted 1:100 in fresh EBB supplemented with 500 µM tryptophan and kynurenine or kynurenic acid. At the start (0h) and at termination of the experiment (24h) 250 µL culture was transferred to a new tube and immediately 1 mL cold (–20 °C) methanol was added. The samples were stored at –20 °C until further use.

### Growth experiments, community construction and sampling in a mini-bioreactor

All experiments were conducted in a MiniBio 250 mL bioreactor (Applikon Biotechnology, The Netherlands) with a working volume of 250 mL. Each experiment was performed in 200 mL EBB + 500 µM tryptophan and, if indicated 100 µM kynurenine. To prevent foam accumulation 0.1 % (v/v) silicon antifoam emulsion (Carl Roth, Germany) was added. The temperature, dissolved oxygen and pH were monitored online. The adaptive my-Control system (Applikon Biotechnology, The Netherlands) was used to control process parameters. Setpoints of 50% of the atmospheric oxygen and 37 °C were applied for dissolved oxygen and the temperature, respectively [39, 40]. The culture was continuously stirred which was controlled with the adaptive my-Control system. The stirrer speed increased linearly with decreasing oxygen below the setpoint up to a maximum of 1200 RPM. The stirrer speed decreased linearly with increasing oxygen above the setpoint down to a minimum of 220 RPM.

To determine kynurenine pathway metabolism of the individually grown bacteria in the bioreactor, a single colony of *P. fluorescens* or 1 mL overnight grown precultures of *E. coli*, *S. salivarius*, *S. thermophilus* or *L. casei* was added to 200 mL EBB supplemented with 500 µM tryptophan and, if indicated, 100 µM kynurenine. At the start (0h) and at termination of the experiment (24h) 250 µL culture was transferred to a new tube and immediately 1 mL cold (–20 °C) methanol was added. The samples were stored at –20 °C until further use.

Community growth was obtained via inoculation of a single colony of *P. fluorescens* MFY63 in 200 mL EBB supplemented with 500 µM tryptophan and, if indicated, 100 µM kynurenine. After 23 hours of growth, 1 mL of combined overnight grown pre-cultures of *E. coli* (1:10000)*, S. Salivarius* (1:100) and *L. casei* (1:100) or *S. thermophilus* (1:100) were added. At 0, 3, 5, 7, and 24 hours after inoculation with the overnight grown pre-cultures a 5 mL sample was obtained. Immediately 250 µL was added to 400 µL NMR buffer (200 mM Na_2_HPO_4_, 44 mM NaH_2_PO_4_, 1 mM TSP, 3 mM NaN_3_ and 20% (v/v) D_2_O), centrifuged at 4 °C, 21130 rcf for 20 min and 550 µL was transferred to a 5 mm NMR tube. For biomass determination, 1.5 mL sample was centrifuged at room temperature for 1 min at 21130 rcf. The supernatant was removed and the wet pellet was weighed. Lastly, 250 µL sample was transferred to a new tube and immediately 1 mL cold (–20 °C) methanol was added. The samples were stored at –20 °C until further use.

### UHPLC-UV

Cell debris was removed from the sample stored at –20 °C in methanol by centrifugation at 4 °C, 21130 rcf for 10 min. The supernatant was transferred to a new tube and the methanol fraction was evaporated using a Savant speed-vacuum dryer (SPD131, Fisher Scientific, Landsmeer, The Netherlands) at 60 °C for 75 min. The dried material was reconstituted to 1 mL with 0.7% perchloric acid. Samples were filtered using a 0.2 µM RC membranes (Phenomenex, Utrecht, The Netherlands) and injected into the UHPLC system (Dionex UltiMate 3000 autosampler; Dionex UltiMate 3000 LPG-3400SD pump, Thermo Fisher Scientific, Waltham, Massachusetts, USA). All samples were analyzed on a C18 column (Kinetex 5 μm, C18 100 Å, 250 × 4.6 mm, Phenomenex, Utrecht, the Netherlands) using a gradient of water/methanol with 0.1% formic acid (0 to 10 min, 95% to 80% H2O; 10 to 20 min, 80% to 5% H2O; 20 to 23 min, 5% H2O; 23 to 31 min, 95% H2O), with a flowrate of 1 mL/min. The column was kept at a temperature of 35 °C. UV-detection at 260 nm was performed with an UV6000LP Detector (Dionex Ultimate 3000 variable wavelength detector, Thermo Fisher Scientific, Waltham, Massachusetts, USA) Data recording and analysis were performed using Chromeleon software (version 6.8 SR13).

### DNA extraction

DNA extraction was performed as previously described [41] from the cell pellets obtained by centrifuging 1.5 mL culture for 1 min at 21130 rcf. The supernatant was removed, the pellets were resuspended in 750 mL lysis buffer (500 mM NaCl, 50 mM Tris-HCl (pH 8.0), 50 mM EDTA, 4 % SDS) and transferred to a 2 mL screw cap tube containing 0.5 g, 0.1 mm zirconia beads and 4, 3 mm glass beads. The cells were disrupted with a mini-bead beater (3X 1 min, with 1 min intervals on ice) and incubated at 95 °C for 15 min. Cell debris was removed by centrifugation at 4 °C, 21130 rcf for 20 min. 500 µL of the supernatant was transferred to a clean tube. 200 µL 10 M ammonium acetate was added, and the samples were incubated for 10 min on ice. The samples were centrifuged at 4 °C, 21130 rcf for 10 min. 600 µL was transferred to a clean tube and 600 µL isopropanol was added. The samples were incubated overnight at –20 °C and afterwards centrifuged at 4 °C, 21130 rcf for 15 min to pellet the nucleic acids. The supernatant was removed and to wash the nucleic acid pellet 700 µL 70 % ethanol was added, the samples were mixed thoroughly and centrifuged at room temperature, 21130 rcf for 5 min. The pellets were completely dried by leaving the tubes open for 60 min at room temperature and were dissolved in 100 µL MilliQ.

### Digital chip-based PCR analysis

Chip-based digital polymerase chain reaction (c-dPCR) was performed with the QuantStudio™ 3D Digital PCR System (User Guide, Catalog Number A29154, Thermo Scientific). Each c-dPCR reaction was prepared in a final volume of 15,5 µL containing 5 µL QuantStudio™ 3D Digital PCR Master Mix v2 (Applied Biosystems), 1 µM end concentration forward and reverse primer, 0,25 µM end concentration probe **(Supplementary table 1, Eurogentec)** and nuclease free water. 1 µL extracted genomic DNA was added to the reaction mix. The DNA concentration was estimated by using the cycle threshold values from the qPCR run. For no-template control 1 µL nuclease free water was added instead of genomic DNA. The positive control was extracted genomic DNA from pure cultures. 14,5 µL was loaded into the QuantStudio™ 3D Digital 20K Chip v2 using the QuantStudio™ 3D Digital PCR Chip Loader. The amplification reactions were performed in the QuantStudio™ 3D Digital PCR System. The PCR conditions were: 10 minutes at 96 °C for DNA polymerase activation, followed by 40 two-step cycles of 2 minutes at 60 °C, 30 seconds at 98 °C and a final extension for 2 minutes at 60 °C and an infinite 10°C hold until the chips were read. After DNA amplification the chips were transferred to the QuantStudio™ 3D Digital PCR Chip reader for imaging. The end-point fluorescence data were collected and analyzed using the QuantStudio™ 3D AnalysisSuite cloud software (version 3.1.6-PRC-build18). Droplets were considered positive when the fluorescence signal was above the threshold. The threshold was set to the positive control DNA and no-template control. The number of positive and negative reactions were counted and using Poisson statistics to measure absolute copies per µL.

## ^1^H NMR spectroscopy and data processing

All ^1^H-NMR spectra were recorded using a Bruker 600 MHz AVANCE II spectrometer equipped with a 5 mm triple resonance inverse cryoprobe and a z-gradient system. The temperature of the samples was maintained at 25 °C during measurement. Prior to data acquisition, tuning and matching of the probe head followed by shimming and proton pulse calibration were performed automatically for each sample. One-dimensional (1D) ^1^H-NMR spectra were recorded using the first increment of a NOESY pulse sequence with presaturation (*γ*B_1_ = 50 Hz) for water suppression during a relaxation delay of 4 s and a mixing time of 10 ms. 256 scans of 65,536 points covering 13,658 Hz were recorded and zero filled to 65,536 complex points prior to Fourier transformation, an exponential window function was applied with a line-broadening factor of 1.0 Hz. The spectra were phase and baseline corrected and referenced to the internal standard (TSP; *δ* 0.0 ppm), using the MestReNova software (v.12.0.0-20080, Mesterlab Research). With the same software, spectral alignment was performed selecting manually areas and applying a linear filling method of the missing values, after the exclusion of the water signal and the surrounding empty area (4.50-5.30 ppm). Spectral binning followed from –0.50 to 9.00 ppm with an equal size binning step of 0.005 ppm. Noise removal was performed by averaging each integrated bin separately and removing the bins with an average below 100. Before ordination analysis bins belonging to TSP (–0.50 – 0.72 ppm), ethanol (1.170 – 1.220 ppm and 3.640 – 3.690 ppm) and a pH sensitive area (7.850 – 8.190 ppm) were removed. The annotation of the bins was performed with the Chenomx Profiler software (Chenomx NMR Suite 8.6 and Chenomx 600 MHz, version 11) and the HMDB database 5.0 (http://www.hmdb.ca).

### Statistical and network analysis

All analysis of variance (ANOVA) were performed using GraphPad Prism 7.0. When a comparison resulted in a statistically significant difference (*P* < 0.05), multiple comparisons testing was performed by controlling the false discovery rate (FDR) according to the Benjamini-Hochberg method (α < 0.05). The R package MixOmics was used for ordination and multivariate statistical analysis of the ^1^H-NMR spectra [42]. The dynamic profile comparison using Kendall’s τ correlation with the BH (α < 0.05) was performed using the R package psych. In CytoScape 3.9.1 the plugin CoNet [43] was used for network construction. The network properties were obtained using the network analyzer tool. To analyze the PLS-DA separation, the accuracy, R2, Q2 and *P*-value of a permutation test performed with the separation distance statistic and 1000 permutations are calculated using MetaboAnalyst 5.0 [44].

## Results

### Kynurenine pathway metabolism and pH changes in a three species synthetic community

To investigate the metabolic network changes resulting from probiotic supplementation, it is critical to carefully select bacterial members for the community. These members should be isolated from the normal small intestinal microbiota and capable of metabolizing key proximal small intestinal metabolites such as, acetate and lactate [7]. Additionally, the kynurenine pathway should be affected by the community. In this study, *Streptococcus salivarius* and two bacteria capable of affecting the kynurenine pathway, namely *Pseudomonas fluorescens* and *Escherichia coli* [33, 45–47], were selected. Two commonly used probiotic species, which were previously shown to be active in the small intestine, *Streptococcus thermophilus* W69 [48, 49] and *Lactobacillus casei* W56 [50], were added separately to distinguish between general and species-specific effects of probiotic supplementation. Before constructing the community, each species was separately grown in flasks with 500 µM tryptophan and kynurenine or KYNA to investigate the ability of the various community strains to metabolize kynurenine pathway metabolites. After 24 hours of incubation, kynurenine pathway metabolism was observed for *P. fluorescens* and *E. coli*, but not for *S. salivarius*, *S. thermophilus* and *L. casei* **(Supplementary figure 1)**. To confirm the kynurenine pathway metabolism in conditions resembling the upper small intestine, each strain is separately grown in a MiniBio 250 mL bioreactor (Applikon Biotechnology, The Netherlands) at 37 °C with controlled pO2 (50% of the atmospheric air) as previously described [39, 40] with and without 100 µM kynurenine. After 24 hours of incubation, the results revealed that *E. coli* effectively metabolized tryptophan and converted kynurenine into KYNA. Conversely, *P. fluorescens* produced only a small amount of kynurenine from tryptophan. In contrast, *S. salivarius*, *S. thermophilus*, and *L. casei* did not metabolize either tryptophan or kynurenine **(Supplementary figure 2A)**. These findings confirm the differential metabolic capabilities of bacterial species in relation to kynurenine pathway metabolism. Concurrently, the pH dynamics of each strain cultured in the presence or absence of kynurenine were assessed. *P. fluorescens* did not exhibit any noticeably pH changes during the course of the experiment, whereas *E. coli*, *S. salivarius* and *S. thermophilus* demonstrated a decrease in pH after 3.5h of growth. *L. casei* reduced the pH after 12h of growth. Among all bacteria, only *E. coli* displayed an increase in pH after 6h. Comparable pH profiles were observed for all bacteria, except for *E. coli* when incubated with 100 µM kynurenine. Co-culturing *E. coli* with kynurenine resulted in an earlier and more pronounced decline in pH **(Supplementary figure 2B)**.

The synthetic community was then grown in the bioreactor under the same conditions stated above. *P. fluorescens* was initially cultured in the bioreactor for a period of 23h. Thereafter, *S. salivarius*, *E. coli* and one of the probiotic species were added to the bioreactor to create a bacterial community. Furthermore, we supplemented each community with 100 µM kynurenine to simulate increased activity of the small intestinal kynurenine pathway. The small intestine is known to undergo rapid changes in response to food ingestion, necessitating the need for the bacterial community to adapt [16, 51]. Moreover, substantial metabolism was observed in individually grown bacteria after 3.5h of growth, as indicated by the pH profile **(Supplementary figure 2B).** Therefore, we collected samples from the community at 0, 3, 5, 7 and 24h after community growth and analyzed the biomass, absolute species distribution using c-dPCR, metabolic environment using ^1^H-NMR, and kynurenine metabolism using HPLC-UV **(Figure 1A)**. Our results showed that both kynurenine (between 3 and 7h) and KYNA (between 7 and 24h) were produced in the control and probiotic-supplemented communities, with kynurenine addition increasing KYNA production in all communities **(Supplementary figure 3)**. However, we observed that the addition of kynurenine together with *L. casei* increased the levels of kynurenine after 5h of growth when compared to the control and *S. thermophilus* supplemented community in the presence of kynurenine and after 7h of growth only in the *S. thermophilus* supplemented community in the presence of kynurenine subsequently leading to increased levels of KYNA after 7 and 24h of growth compared to the control and *S. thermophilus* supplemented communities **(Figure 1B)**. Moreover, the pH declined sharply after 3h of community growth, followed by a linear increase and a plateau after 10h of growth **(Figure 1C)**, and the addition of kynurenine did not affect the lowest pH and the time it was reached (ordinary two-way ANOVA; F(1,12) = 1.78, *P* = 0.21 and F(1,12) =0.76, *P* = 0.4 respectively) **(Table 1)**. Probiotic supplementation altered both the lowest pH and the time it was reached (ordinary two-way ANOVA; F(2,12) = 6.85, *P* = 0.01 and F(2,12) = 4.0, *P* = 0.047) **(Table 1)**, mostly attributed to differences between *L. casei* and *S. thermophilus* supplementation **(Figure 1C, Supplementary figure 4)**, indicating a probiotic specific effect.

**Figure 1:**
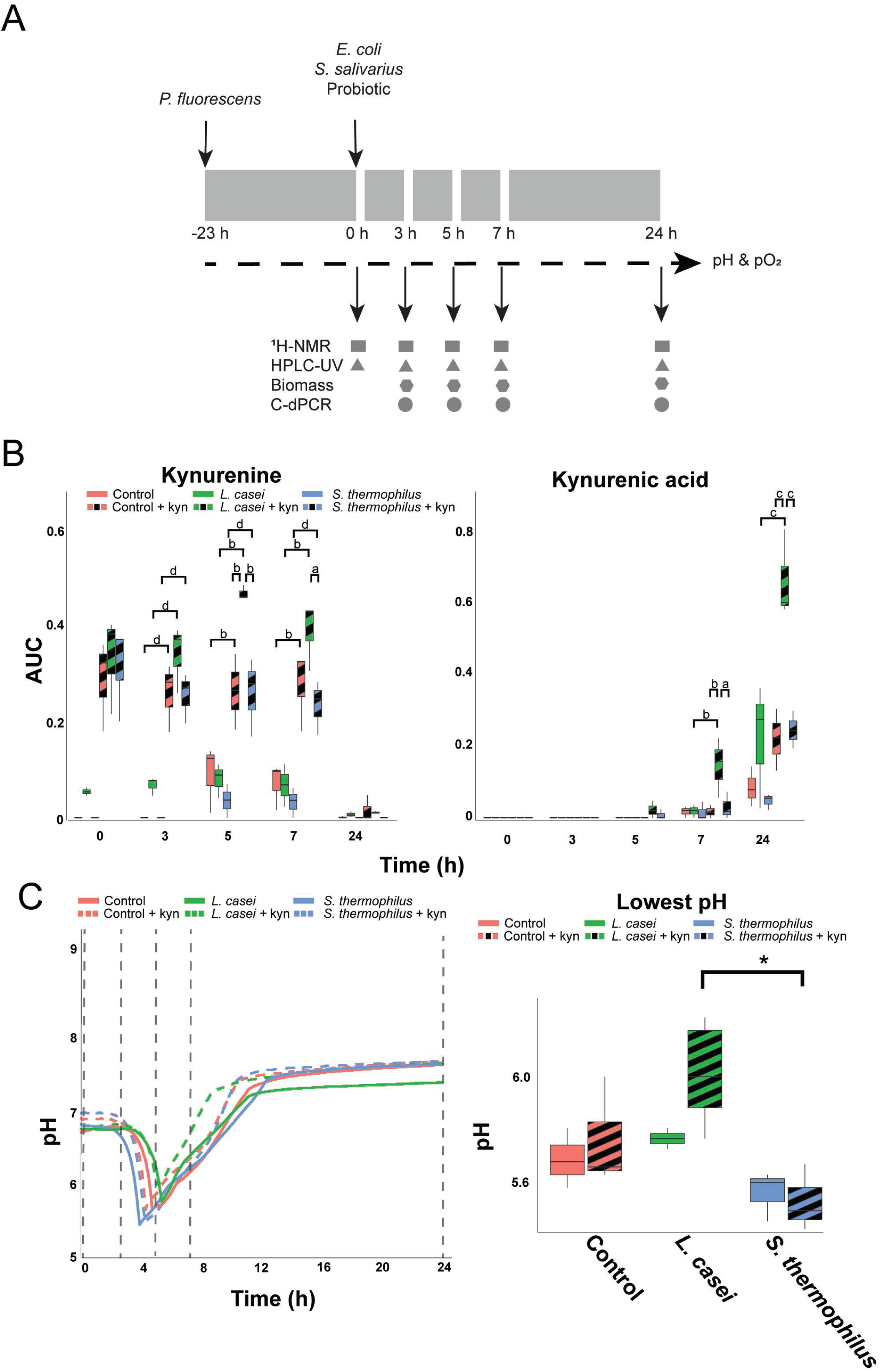
*Lactobacillus casei* supplementation increases kynurenine pathway metabolism in a synthetic community. A) Overview of the construction of the community and sampling procedure. *P. fluorescens* was inoculated 23h prior to inoculating other species with probiotics. PH and pO_2_ (indicated with dashed line) were continuously measured. Black arrows indicate sampling points (0, 3, 5, 7 and 24h) for obtaining metabolomics, compositional and biomass data. All experiments were performed in triplicates. B) Boxplots showing the area under the curve (AUC) for kynurenine and kynurenic acid as measured from the HPLC-UV chromatograms normalized to the AUC of the initial tryptophan peak obtained from –23h. Significant comparisons are depicted with letters (a-d): a: q < 0.05, b: q < 0.01, c: q < 0.001 and d: q < 0.0001 as obtained with an ordinary two-way ANOVA and multiple comparisons testing by controlling the false discovery rate according to the Benjamini-Hochberg procedure. C) pH profile for control, *S. thermophilus* supplemented and *L. casei* supplemented communities with/without kynurenine. Sampling points indicated by dashed lines. Boxplot indicates the lowest pH value per condition. Significant differences are indicated with asterisks; *, *q* < 0.05, as obtained with an ordinary one-way ANOVA and multiple comparison testing by controlling the false discovery rate according to the Benjamini-Hochberg procedure.

**Table 1:**
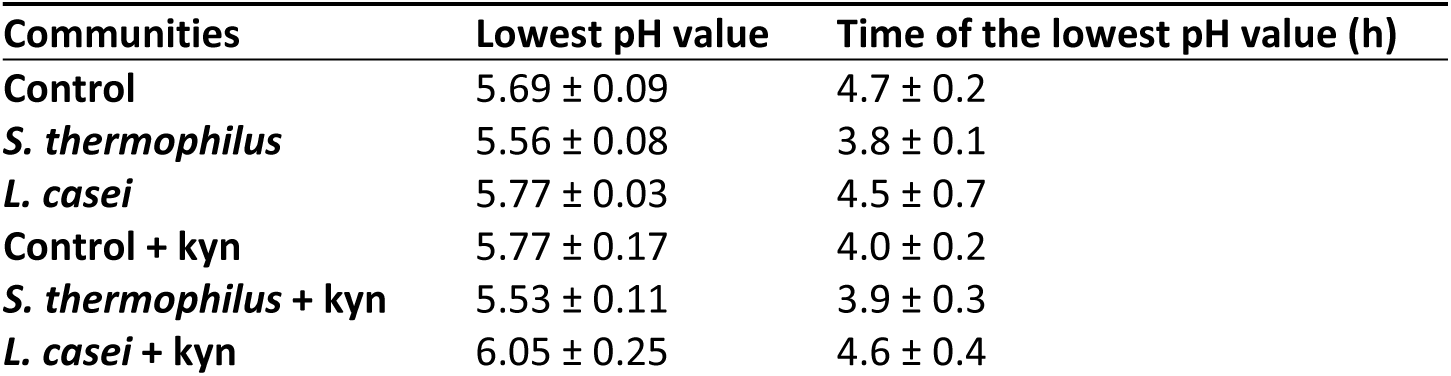
properties of the pH profile of each community. The mean ± SD of the lowest pH value and time the lowest pH value was reached for all communities with and without kynurenine (kyn) supplementation.

| Communities | Lowest pH value | Time of the lowest pH value (h) |
| --- | --- | --- |
| **Control** | 5.69 ± 0.09 | 4.7 ± 0.2 |
| ***S. thermophilus*** | 5.56 ± 0.08 | 3.8 ± 0.1 |
| ***L. casei*** | 5.77 ± 0.03 | 4.5 ± 0.7 |
| **Control + kyn** | 5.77 ± 0.17 | 4.0 ± 0.2 |
| ***S. thermophilus* + kyn** | 5.53 ± 0.11 | 3.9 ± 0.3 |
| ***L. casei* + kyn** | 6.05 ± 0.25 | 4.6 ± 0.4 |

Overall, the addition of kynurenine together with *L. casei* increased the production of kynurenine and subsequently KYNA compared to the control and *S. thermophilus* supplemented communities. Probiotic supplementation altered both the lowest pH and the time it was reached, mostly attributed to differences between *L. casei* and *S. thermophilus* supplementation, indicating a probiotic-specific effect. These findings suggest that probiotics, particularly *L. casei*, may influence the production of kynurenine pathway metabolites in the small intestine, which could have important implications for human health.

### Effect of kynurenine and probiotic supplementation on biomass and cell counts in a simplified bacterial community

Bacteria can gain a competitive advantage in a community through metabolizing compounds present in the environment, inhibiting the growth of competitors, or stimulating other bacteria to secrete beneficial metabolites [52]. Therefore, we hypothesized that the supplementation of a probiotic species or kynurenine, which was only metabolized by *P. fluorescens* and *E. coli* **(Supplementary figure 2A)**, may alter the composition of the simplified community. The total biomass and the community composition were compared **(Figure 1A)** to investigate the effects of kynurenine and probiotic supplementation on community growth. Kynurenine addition did not affect the biomass, while probiotic addition altered total biomass production after 7h of growth **(Supplementary data 1).** Specifically, the addition of *L. casei* resulted in an increase in biomass in the absence of kynurenine, while the introduction of *S. thermophilus* in the presence of kynurenine led to a reduction in the total biomass **(Supplementary figure 5)**.

Next, the total cell count and cell count of specific species in the community were monitored using c-dPCR. Kynurenine supplementation did not affect the total cell count or the cell count of any individual species **(Supplementary data 2).** Furthermore, probiotic supplementation did not alter the cell count of *P. fluorescens* **(Figure 2, Supplementary data 2)**, but probiotic supplementation altered the total cell count and the cell count of *E. coli* (after 3 and 5h of growth, although multiple comparison testing did not identify specific alterations), and of *S. salivarius* (after 5 and 7h of growth) **(Figure 2, supplementary data 2)**, which coincides with the decrease in pH **(Figure 1C)** and with the growth of *S. thermophilus* **(Figure 2).** In contrast, the cell counts of *L. casei* continuously increased over time **(Figure 2)**. Cell counts of the probiotic species were not affected by kynurenine supplementation at any sampling point **(Figure 2)**. Specifically, *L. casei* addition inhibited the growth of *S. salivarius* by altering the community metabolism. These results suggest that the supplementation of certain compounds, exemplified here by kynurenine, or probiotics, or a combination of both, can alter the growth and composition of bacterial communities, which could have implications for the design of probiotic therapies.

**Figure 2:**
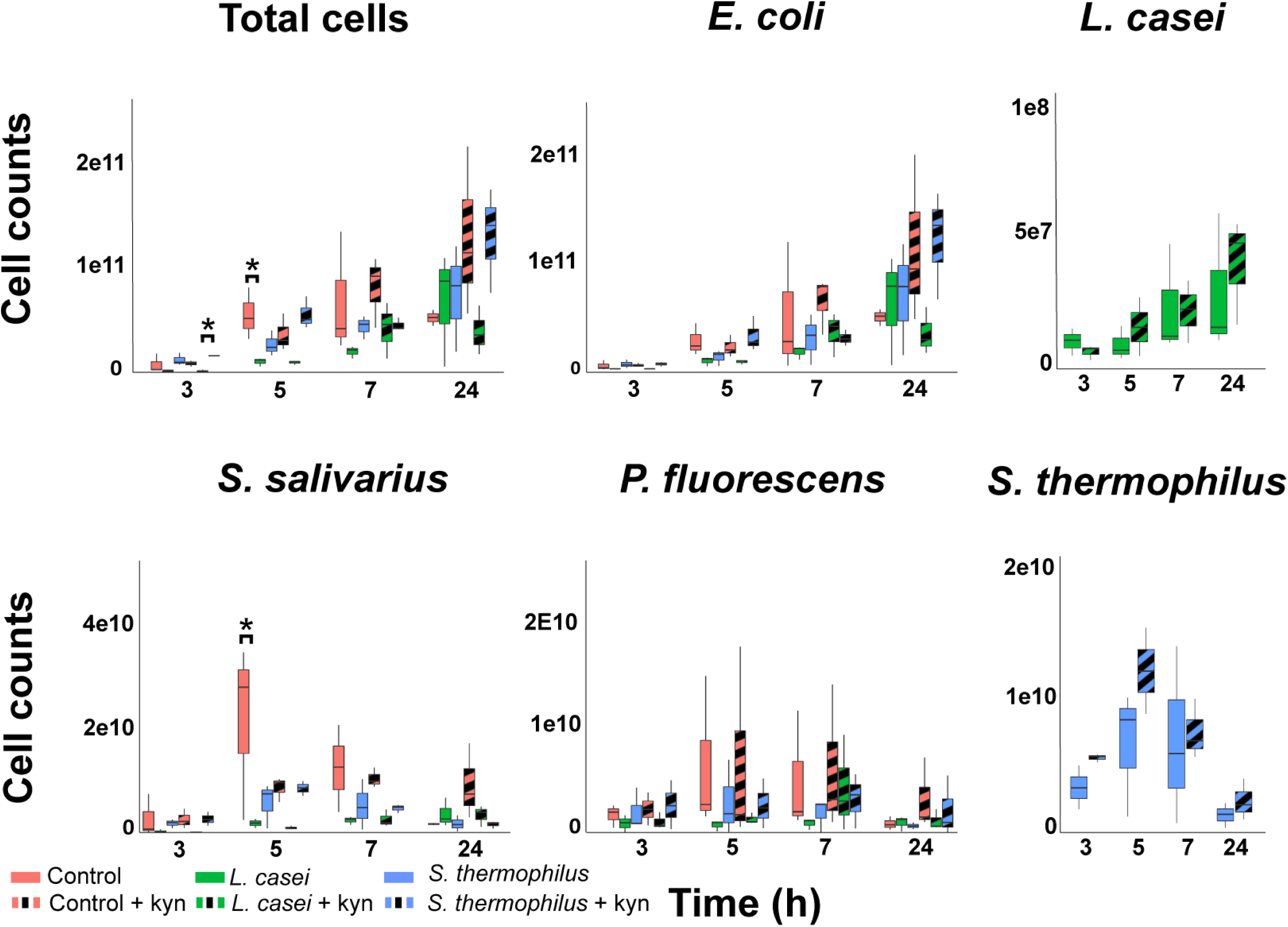
Supplementation of *Lactobacillus casei* reduces the growth of *Streptococcus salivarius* in a simplified community. Boxplots of total cell count and cell count per species of a simplified community (*E. coli*, *S. salivarius* and *P. fluorescens*) determined after 3, 5, 7 and 24h of growth. Cell counts of the communities grown with the supplemented probiotic species *L. casei* or *S. thermophilus* are depicted in different colors. Control or probiotic supplemented communities grown with 100 µM kynurenine are shown with stripes. Statistical differences between communities were determined by ordinary two way ANOVA approach per timepoint, and significant differences (q < 0.05) after multiple comparisons testing by controlling the false discovery rate according to the Benjamini-Hochberg procedure are indicated with asterisks. Probiotic addition altered the cell count of *E. coli* after 3 and 5h of growth, and the cell count of *S. salivarius* after 7h of growth (ordinary two-way ANOVA: F(2,12) = 4.43 and P = 0.036, F(2,12) = 4.15, P = 0.043 and F(2,12) = 7.62, P = 0.007 respectively), but multiple comparisons testing did not yield any significant differences. Cell counts of added probiotic species grown with and without kynurenine per timepoint were compared by performing an unpaired t test.

### Metabolic network analysis of probiotic supplementation and kynurenine metabolism in the synthetic bacterial community

In order to elucidate the metabolic changes that are associated with alterations in pH, kynurenine metabolism and cell count, we employed ^1^H-NMR to analyze the supernatant **(Figure 1A).** Following spectral binning and processing, a total of 943 bins were subjected to principal component analysis (PCA). Our results revealed time-dependent clustering of the samples **(Figure 3A)**, whereby the most significant contributors to the observed separation were lactate (bin 366, 368 and 369), acetate (bin 485 and 486).

**Figure 3:**
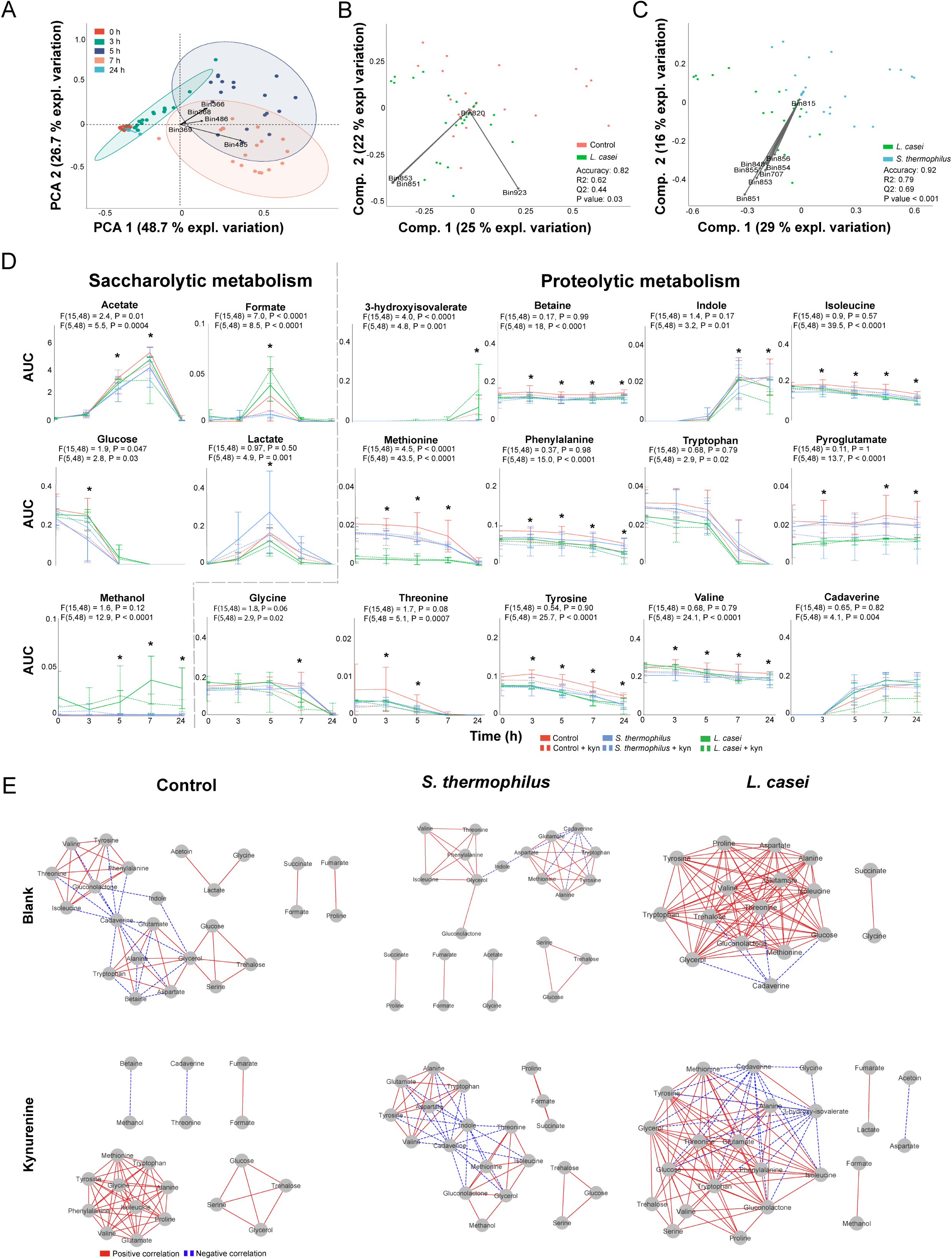
Probiotic and kynurenine supplementation alter the metabolism of a simplified community in a specific manner. A) PCA with binned ^1^H-NMR spectra as input showing separation of metabolites with lactate (bin 366,368 and 369), acetate (bin 485 and 486) being the most distinguishing factors (indicated with arrows). The ellipses represent the 95 % confidence interval. B) PLS-DA with binned ^1^H-NMR spectra of control and *L. casei* supplemented communities, indicating lactate (bin 923), glucose (bin 851 and 853), and threonine (bin 820) as contributing factors. C) PLS-DA with binned ^1^H-NMR spectra of *S. thermophilus* and *L. casei* supplemented communities indicating glucose (bin 848, 851 and 853 – 856), glycerol, (bin 815) and cadaverine (bin 707) as contributing factors. Statistical parameter accuracy, R2 and Q2 and *P-*values are included. The *P*-value is the results of a permutation test performed with the separation distance statistic and 1000 permutations. All parameters were calculated using MetaboAnalyst 5.0. D) Dynamic profiles of the significantly different saccharolytic and proteolytic metabolites with repeated measure two-way ANOVA showing significant differences. The error bars indicate the standard deviation. The timepoints where a significant result of multiple comparison testing by controlling the false discovery rate according to the Benjamini-Hochberg procedure is obtained are indicated with an asterisk; *: q < 0.05, detailed results of the multiple comparison tests can be found in the supplementary data 3. E) Dynamic correlation-based networks of all measured metabolites in the communities showing positive and negative correlations via red solid and blue dashed edges, respectively. Nodes represent metabolites. Edges represents correlations between the two connected nodes, obtained when 2/3 methods give a positive results; Kendall’s –0.9 > τ > 0.9, Spearman’s –0.9 > ρ > 0.9. Brown’s randomization method with 1000 iterations had a Benjamini-Hochberg corrected *P-*value < 0.05. The calculation of the edges was performed using the Cytoscape plugin CoNet.

To determine whether the dynamic metabolic profiles of the community were altered by probiotic or kynurenine supplementation, we conducted partial least squares – discriminant analysis (PLS-DA) with cross validation (CV) and a permutation test was performed with the 943 bins as input. Our analysis revealed no significant differentiation between communities with and without kynurenine supplementation, nor between control and *S. thermophilus*-supplemented communities **(Supplementary figure 6A and 6B)**. However, we observed significant separation in communities supplemented with *L. casei,* with lactate (bin 923), glucose (bin 851 and 853) and threonine (bin 820) contributing most to the separation from the control **(Figure 3B)**. Additionally, glucose (bin 848, 851 and 853 – 856), glycerol (bin 815) and cadaverine (bin 707) were the major contributors to the differentiation between *L. casei* and *S. thermophilus* supplemented communities **(Figure 3C)**.

To demonstrate the metabolic contributions responsible for the variance obtained from the entire spectral dataset, the chemical library of Chenomx profiler software and the online human metabolome database version 5.0 were utilized. We identified 30 metabolites, and subjected the area under the curve (AUC) of a representative peak for each metabolite to ordination analysis. The results of this approach were similar to the ordination analyses conducted with the bins. PCA revealed temporal clustering of the samples and PLS-DA revealed significant separation of the *L. casei* supplemented community from the control and *S. thermophilus* supplemented communities **(Supplementary figure 7A, B and C)**. Therefore, we postulated that the AUC values of the identified set of metabolites may serve as a surrogate for the full spectra dataset and could be employed for subsequent analysis.

Given that the metabolic network of a community is highly interconnected, a substantial alteration in certain metabolites can lead to multiple smaller changes in related metabolites that might not be detectable by ordination analysis **(Figure 3A, B and C)**. Similarly, minor effects of kynurenine supplementation on metabolism may go undetected via ordination analysis. To investigate smaller metabolic changes induced by kynurenine or probiotic addition to the synthetic community, we compared the dynamic metabolic profiles between experiments, which revealed similarities **(Supplementary figure 8)** and differences in the dynamics of saccharolytic and proteolytic community metabolism **(Figure 3D, Supplementary data 3)**. Of note, we observed increased lactate production in the *S. thermophilus* supplemented community, which could potentially account for the observed pH variation **(Figure 1C)**.

To explore the coordinate behavior of metabolites and network topology features across the tested conditions [53], we constructed dynamic correlation-based networks using the Cytoscape plugin CoNet [43], whereby the nodes represent the metabolites and the edges represent correlations between the metabolic profiles per condition. Without kynurenine, *S. thermophilus* supplementation decreased the number of edges and the clustering coefficient, while the metabolite with the highest number of edges was aspartate (7), compared to cadaverine (12) in the control network **(Figure 3E**, **table 2)**. *L. casei* supplementation had a more drastic effect on the network, increasing the number of edges, clustering coefficient and network density substantially **(Figure 3E**, **table 2)**. Kynurenine supplementation reduced the number of edges, including indole, present in the control community network, and reduced the number of edges of cadaverine from 12 to 1. Consequently, cadaverine and indole no longer connected two major hubs of connected metabolites compared to control community. The clustering coefficient and network density of the control community supplemented with kynurenine was 1, due to the connections between all the metabolites with all the other metabolites in each hub **(Figure 3E**, **table 2)**. Kynurenine supplementation to the *S. thermophilus* and *L. casei* supplemented communities increased the number of edges, clustering coefficient and network density, with indole and cadaverine being the metabolites with the most edges, similar to the control network without kynurenine supplementation **(Figure 3E**, **table 2).** The node degree and betweenness centrality of individual metabolites were also analyzed, showing probiotic-specific effects on the community metabolism as well as an effect of kynurenine supplementation on cadaverine metabolism **(Supplementary data 4)**. Together the network analysis revealed probiotic-specific effects on the community metabolism and an effect of kynurenine supplementation on clustering coefficient, network density and node parameters of the metabolic networks, mainly due to alterations in cadaverine utilization.

**Table 2:** Properties of the dynamic correlation-based metabolic networks for each community. The number of nodes, edges, clustering coefficient, network density and the nodes with the most edges of the networks depicted in figure 3E for the communities with probiotic or kynurenine (kyn) supplementation.

| Communities | Nodes | Edges | Clustering coefficient | Network density | Node with the most edges (edges) |
| --- | --- | --- | --- | --- | --- |
| **Control** | 24 | 54 | 0.857 | 0.368 | Cadaverine (12) |
| ***S. thermophilus*** | 23 | 40 | 0.794 | 0.374 | Aspartate (7) |
| ***L. casei*** | 17 | 94 | 0.935 | 0.886 | Gluconolactone/gluconate, glycerol (14) |
| ***L. casei* + kyn** | 24 | 103 | 0.892 | 0.654 | Glucose, tryptophan, glycerol (16) |
| ***S. thermophilus* + kyn** | 20 | 56 | 0.901 | 0.549 | Indole, cadaverine (12) |
| **Control + kyn** | 20 | 54 | 1 | 1 | Tryptophan, phenylalanine, alanine, methionine, glutamate, proline, isoleucine, glycine, valine, tyrosine (9) |

## Discussion

The present study investigated the impact of probiotic supplementation on metabolism of the small intestinal microbiota. To achieve this, we constructed a synthetic community consisting of three gut isolates that mimicked the microbial community of the upper small intestine. Specifically, we examined the metabolic impact of probiotic supplementation on both general metabolism and the kynurenine pathway, which has been implicated in the modulation of immune responses and neuroinflammation. Our findings shed light on the probiotic-specific mechanisms of action that drive colonization resistance in the small intestine and suggest that the kynurenine pathway may be a promising target for future probiotic-based therapies aimed at promoting intestinal health.

Probiotics are known to exert their effects through competitive exclusion often by reducing the pH of the gut milieu through increased lactate production [54]. In our study, supplementation with *S. thermophilus* led to increased lactate production, resulting in a reduction in pH, and accompanied by a decrease in formate and acetate levels **(Figure 3D and 1C)**. However, this supplementation did not result in significant changes to the composition of the community **(Figure 2)** or the dynamic metabolic network **(Figure 3E)**. Conversely, when *L. casei* was supplemented, we observed a reduction in the growth of *S. salivarius*, which was not accompanied by increased lactate production **(Figure 2, 3D)**. Other studies have also reported beneficial effects of *L. casei* supplementation independent of acidification or lactate production [55]. These results suggest that *L. casei* supplementation alters the metabolic environment, improving metabolic utilization and thereby, possibly, enhancing the resistance of the community to perturbations **(Figure 3D and E).** This finding is consistent with previous studies showing a positive association between habitual *L. casei* Shirota intake and long-term compositional stability of the fecal microbiota [56].

The concept of direct colonization resistance refers to the ability of the resident microbiota to resist the colonization and expansion of invading species [57]. However, decreased diversity can lead to unoccupied ecological niches that can be occupied by invading species potentially harmful to host health [58]. Probiotics have been proposed as a means to increase colonization resistance by filling unoccupied niches before pathogenic species can colonize [59]. However, current probiotic interventions do not take into account the functional capacity of probiotics to occupy specific niches in the gut [60], which may explain the variable effectiveness of probiotics [61]. To address this, synthetic communities are constructed [22, 62–65], but the lack of molecular insights has hampered the development of probiotic therapies. The genetic capacity of *L. casei* in our synthetic community of three species made it more functionally diverse **(Figure 3D)** compared to the three initial community members and compared to *S. thermophilus*, which resembles *S. salivarius*. Thus, our results suggest several hypotheses on how the tested species mutually influence each other and consequently impact the kynurenine pathway **(Figure 4)**. These hypotheses can be tested in future studies and potentially expanded to include other species.; First, *L. casei* increased the metabolic utilization and, possibly, the resistance of the community to perturbations **(Figure 3E**, **table 2)**, by occupying an empty ecological niche, whereas *S. thermophilus* had to compete for the ecological niche occupied by *S. salivarius.* However, the ability to occupy a niche is determined by the functional capability [66] and the environment [67]. Therefore, selecting probiotic strains with a wide range of functional capacities and constructing a diverse multispecies probiotic formula could be a promising approach for developing effective probiotic therapies [18, 68].

**Figure 4:**
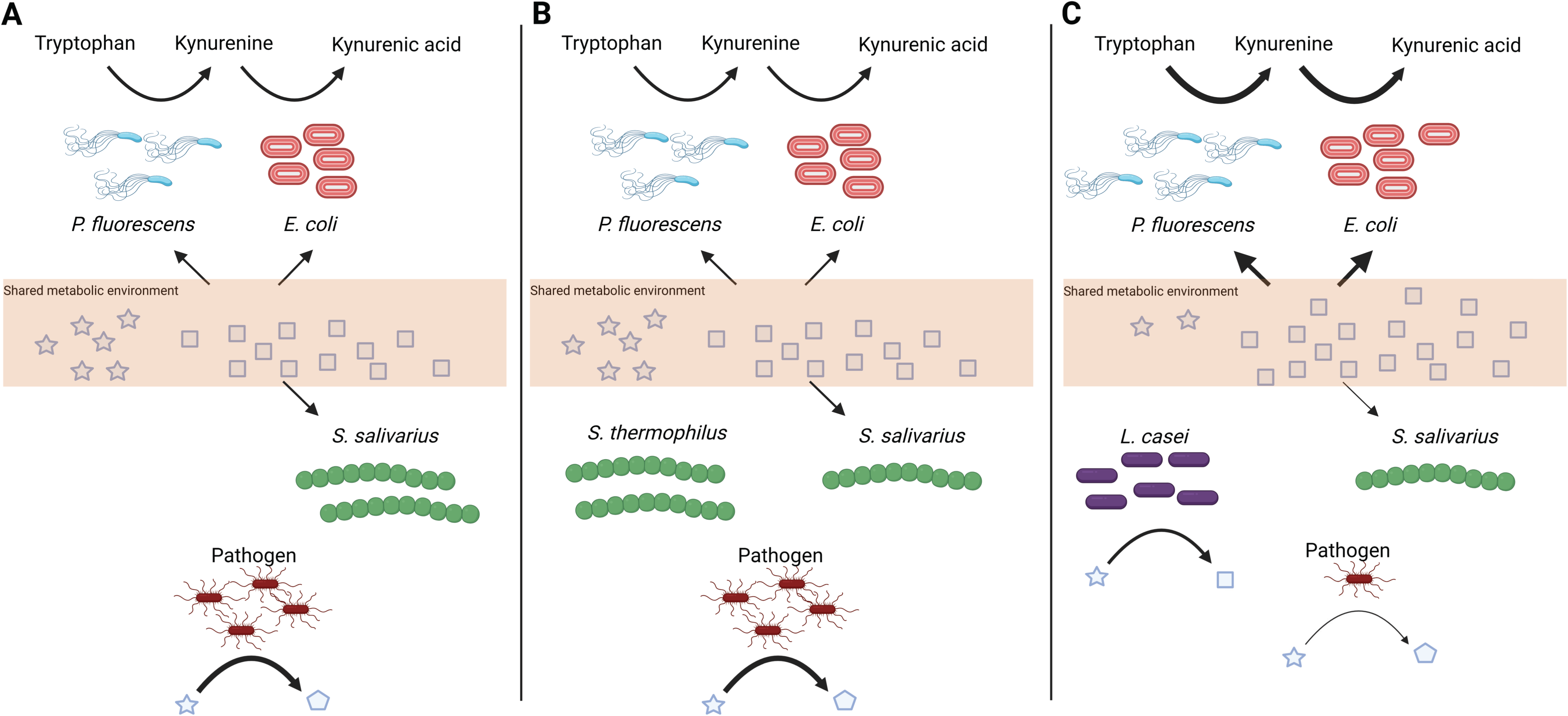
Possible hypotheses of how our tested species affect each other and thereby the kynurenine pathway. A) Microbial community composed of *P. fluorescens*, *E. coli*, and *S. salivarius* can metabolize certain metabolites present in their shared metabolic environment (depicted in squares), while being unable to metabolize others (depicted in stars). If an invading pathogen can metabolize the unused metabolites, it may colonize and produce metabolites with a negative impact on host health (depicted in pentagons). Probiotic intervention may prevent the colonization of invading pathogens, but only if the functional profile of the probiotic species complements the functional capacity of the existing microbiota. In our tested species, B) *S. thermophilus* is functionally similar to *S. salivarius*; thus, the invading pathogen would still be able to colonize and negatively affect host health. On the contrary, C) *L. casei* is functionally different from the existing microbiota and can metabolize the unused metabolites (stars) into metabolites that the existing microbiota can utilize. As a result, an invading pathogen cannot occupy the ecological niche, preventing colonization and positively affecting host health. Additionally, the increased production of metabolites by *L. casei* enhances the metabolism of the existing microbiota, consequently increasing the cross-feeding reaction between *P. fluorescens* and *E. coli* regarding the kynurenine pathway, which positively affects the host as well.

Indirect colonization resistance involves improving the gut epithelial barrier or enhancing the hosts immune system [68]. Metabolites produced by the microbiota can interact with the host and improve gut integrity. For example, stimulation of GPR35 by KYNA was shown to enhance mucosal repair and stimulate mucus secretion, which increased the integrity of the gut epithelial barrier [69, 70]. Thus, increasing microbial production of KYNA in the small intestine may indirectly increase colonization resistance. Although no in vivo studies were conducted in our study to confirm improved intestinal barrier integrity due to increased microbial KYNA production resulting from probiotic addition, we show that in the *L. casei* supplemented community, *E. coli* had a reduced impact on the metabolic environment since the node degree and betweenness centrality of the *E. coli* produced metabolites cadaverine [71] and indole [72] were lower **(Figure 3E)**. In the kynurenine and *S. thermophilus* supplemented community*, E. coli* had an increased impact on the metabolic environment **(Figure 3E),** even though this effect was not translated into increased cell counts for *E. coli* **(Figure 2)**.

Overall, the present study demonstrates that supplementation with different probiotics can lead to distinct alterations in the metabolic environment of the microbiota community, with *L. casei* showing improved metabolic utilization. These findings highlight the importance of selecting probiotic species with diverse genetic capabilities that complement the functional capacity of the resident microbiota or constructing a multispecies formula. Such an approach holds promise for the development of effective probiotic therapies. It emphasizes the need to consider the functional capacity of probiotic species when designing interventions. Furthermore, while the probiotic species may not have directly impacted the kynurenine pathway, they still had an influence on the metabolic profiles through interactions with other species. This highlights the importance for researchers to consider not only the direct effects of probiotic species but also their impacts on the overall community metabolism. Moreover, the present findings suggest that increasing microbial production of metabolites such as KYNA in the small intestine may indirectly improve intestinal barrier integrity and increase colonization resistance. Further in vivo studies are necessary to confirm the effects of probiotic supplementation on intestinal barrier integrity and to develop probiotic therapies aimed at specific ecological niches in the gut.

## Author Contributions

JJ, SEA: Conceptualization, investigation, writing-original draft, and editing; JJ, ACC, KC: data analysis; JJ, ACC: methodology; SvH: Conceptualization, writing review; SEA: funding acquisition. All authors have read and agreed to the published version of the manuscript.

## Acknowledgements

We thank Valeria Mathlener of Winclove, B.V. assisting us with the c-dPCR measurements.

## Data availability

The raw ^1^H-NMR data was deposited in the Metabolights public repository [73] under the accession number: MTBLS7370

