## Supplementary figures for "Metabolic network construction reveals probiotic-specific alterations in the metabolic activity of a synthetic small intestinal community"

^2^ Winclove Probiotics, Amsterdam, the Netherlands

*Present address: Department of Chemistry, Laboratory of Analytical Biochemistry, University of Crete, Heraklion, Greece

^#^ Corresponding author

Sahar El Aidy. Host-microbe Interactions, Groningen Biomolecular Sciences and Biotechnology Institute (GBB), University of Groningen, Nijenborgh 7, 9747 AG Groningen, The Netherlands. P: +31(0)503632201.

E:

**Supplementary Table 1: Primers and probes used for the digital chip-based PCR analysis**

| Strain | Oligo type | Sequence 5’-3’ |
| --- | --- | --- |
| *Escherichia coli* | Forward | TACCTATAGCGGTGCGGAGA |
| *Escherichia coli* | Reverse | TCGCCCTGTTTGTCTTTGGT |
| *Escherichia coli* | Probe | TGTCCCTCACCCGCTTGTTGATCC |
| *Pseudonomas fluorescens* | Forward | TCGGTGCAAAGCTACCAGTT |
| *Pseudonomas fluorescens* | Reverse | ATCAGGCTCTGCAACGACTC |
| *Pseudonomas fluorescens* | Probe | CTCGACCCGCTTCTGGAACGC |
| *Streptococcus salivarius* | Forward | ACAGAAATCACGAAGACTCCAGA |
| *Streptococcus salivarius* | Reverse | CTTCTTACTCCAACGCTTCGA |
| *Streptococcus salivarius* | Probe | AGGTCCTCTATCTTCCTCTATTAGCGC |
| *Streptococcus thermophilus* | Forward | CAAAGGGAAAGTTTCATCTCCAAAA |
| *Streptococcus thermophilus* | Reverse | CTCCAAAATGATCAGCAACTTAAACA |
| *Streprococcus thermophilus* | Probe | ACCTCTCCGCCCTCTCAATTGTATACTGT |
| *Lactobacillus casei* | Forward | AGCTGGACAAGTTCGTTCATGCC |
| *Lactobacillus casei* | Reverse | TTGGCACGATCAGTTGTCGCATAA |
| *Lactobacillus casei* | Probe | TGTTGCGATTGGGATTGGCGGTTC |

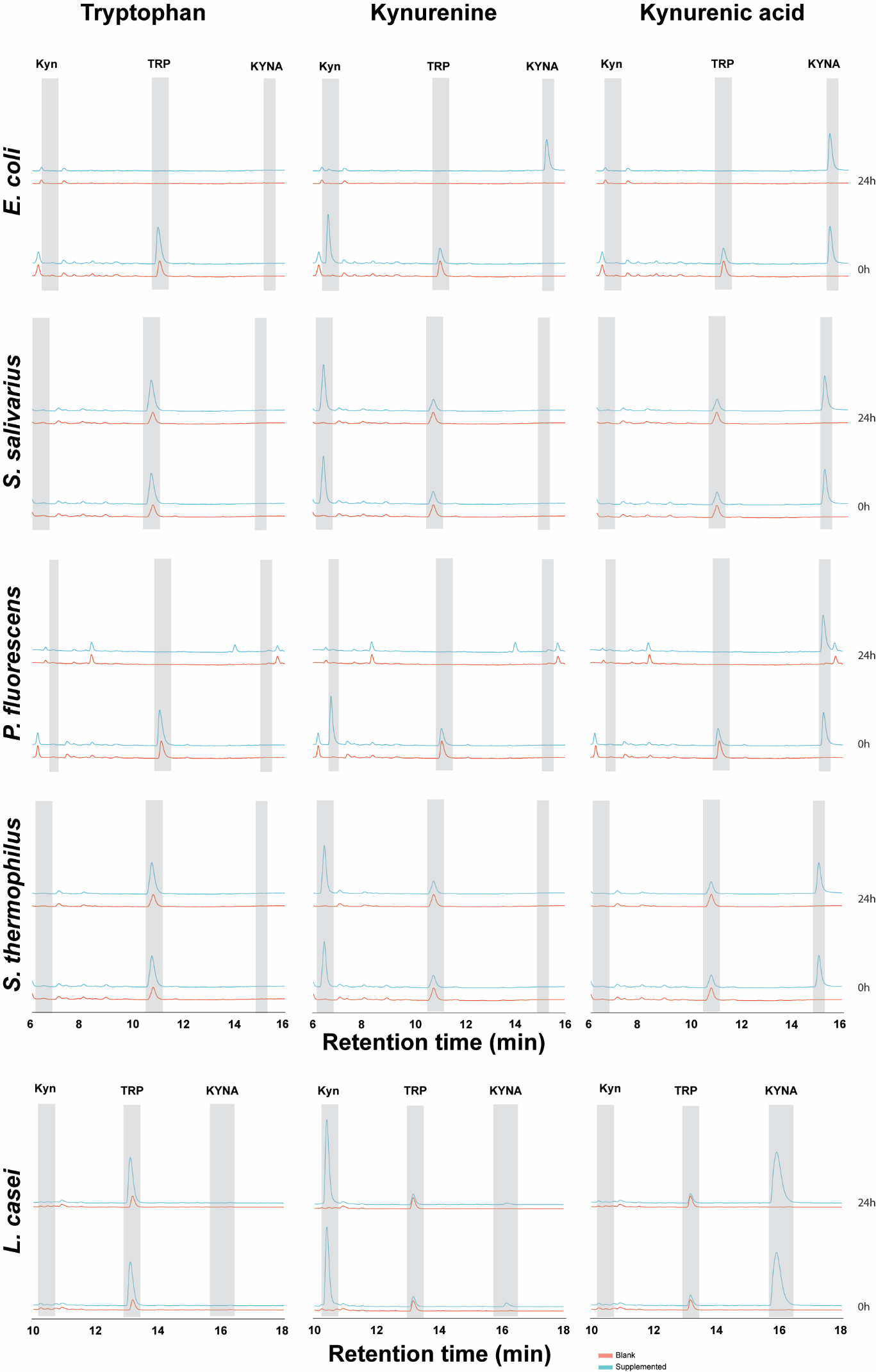

**Supplementary figure 1: kynurenine pathway metabolism of the members of the synthetic community grown separately**

HPLC-UV chromatograms of the supernatant of all members of the synthetic community, *Escherichia coli, Streptococcus salivarius* and *Pseudomonas fluorescens*, and the probiotic species, *Streptococcus thermophilus* and *Lactobacillus casei*. Each bacterium is grown separately for 24h in EBB (Blank) or in EBB supplemented with 500 µM tryptophan (TRP), kynurenine (Kyn) or kynurenic acid (KYNA). The chromatograms depict the supernatant obtained at the start (0h) and end (24h) of the incubation. The retention times of Kyn (6.8min), TRP (11.1min) and KYNA (15.2min) are highlighted with grey bars. The bacteria without supplementation are depicted in red, the bacteria with supplemented kynurenine pathway metabolites are depicted in blue. The supernatant of *L. casei* was analyzed using different HPLC setting, therefore the retention times of Kyn (10.4min), TRP (13.1min) and KYNA (15.9min) are altered.

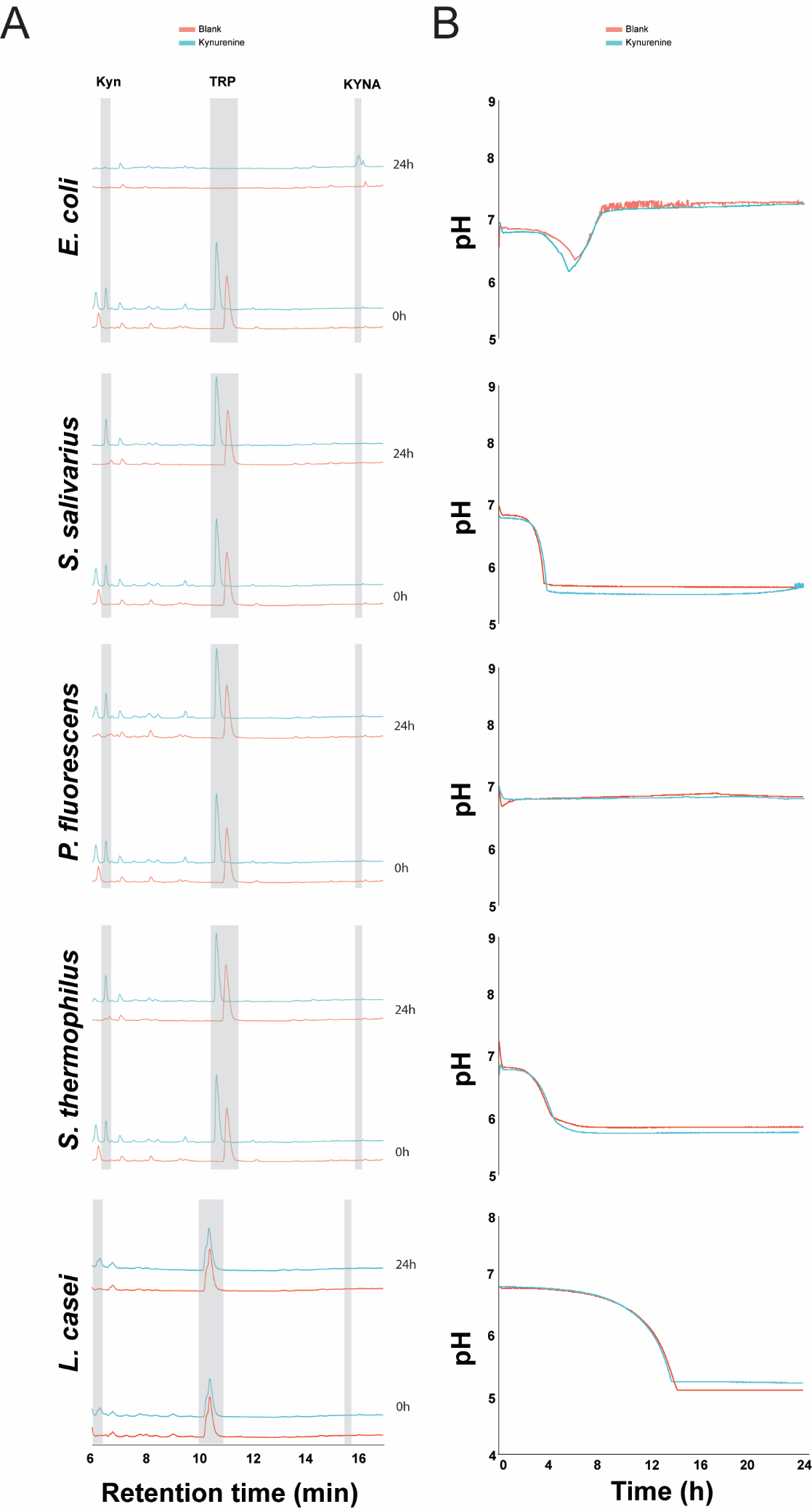

**Supplementary figure 2: kynurenine metabolism and pH of the members of the synthetic community grown separately in a Mini bioreactor**

A) HPLC-UV chromatograms of the supernatant of all members of the synthetic community, *Escherichia coli, Streptococcus salivarius* and *Pseudomoans fluorescens*, and the probiotic species, *Streptococcus thermophilus* and *Lactobacillus casei*. Each bacterium is grown separately for 24h in EBB (Blank) or in EBB supplemented with 100 µM kynurenine (Kyn). The chromatograms depict the supernatant obtained at the start (0h) and end (24h) of the incubation. The retention times of Kyn (6.8min), tryptophan (TRP) (11.1min) and kynurenic acid (KYNA) (15.2min) are highlighted with grey bars. The bacteria without supplementation are depicted in red, the bacteria with supplemented kynurenine pathway metabolites are depicted in blue. The supernatant of *L. casei* was analyzed using a different column of the same type, therefore the retention times are slightly shifted and a shoulder is present at the peaks. B) pH profile measured throughout the incubation experiment described for panel A.

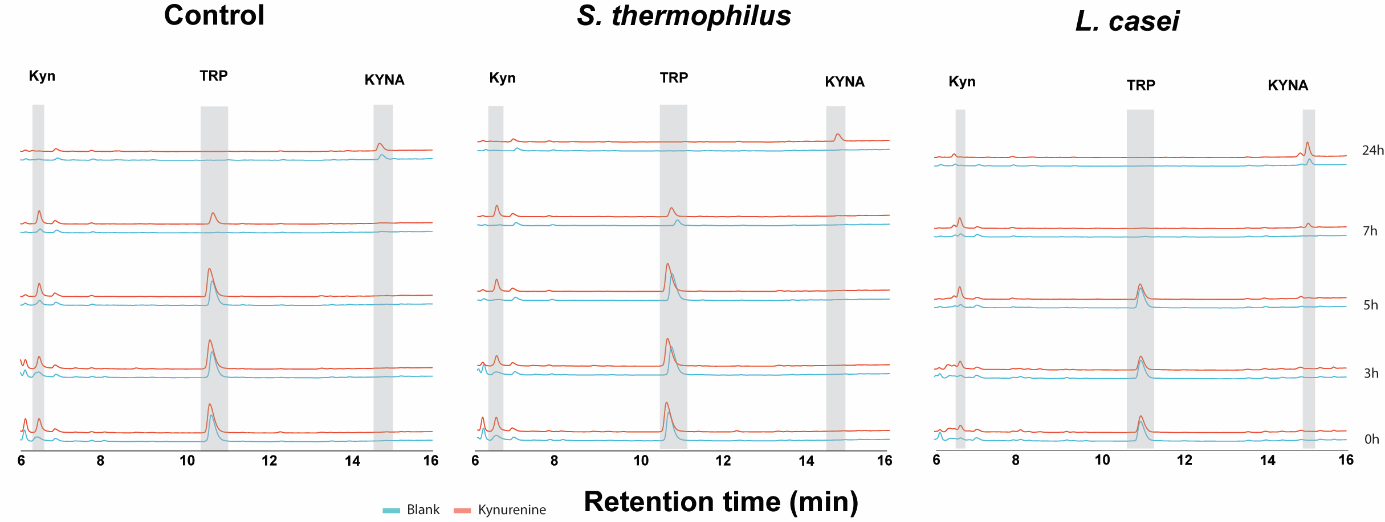

**Supplementary figure 3: representative examples of kynurenine pathway metabolism in the simplified communities**

Representative examples of HPLC-UV chromatograms obtained for each sampling point per community. The blue line indicates the community without kynurenine supplementation, the red line indicates the communities supplemented with 100 µM kynurenine. The retention times of kynurenine (kyn), tryptophan (TRP) and kynurenic acid (KYNA) are indicated with grey boxes.

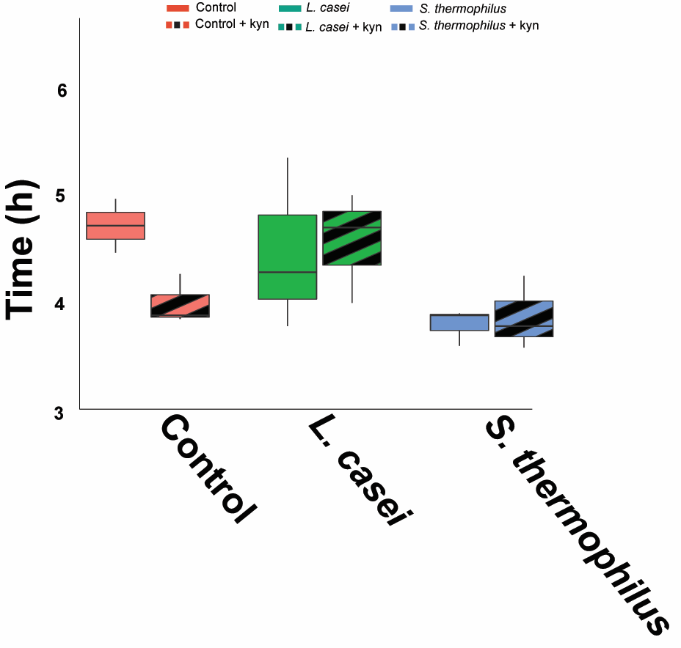

**Supplementary figure 4: the time the lowest pH value was reached in each community**

The boxplots indicate the time the lowest pH value was reached. A significant difference upon probiotic was obtained (ordinary two-way ANOVA: F(2,12) = 4.0, P = 0.047), but multiple comparisons testing by controlling the false discovery rate according to the Benjamini-Hochberg procedure did not result in specific differences between conditions.

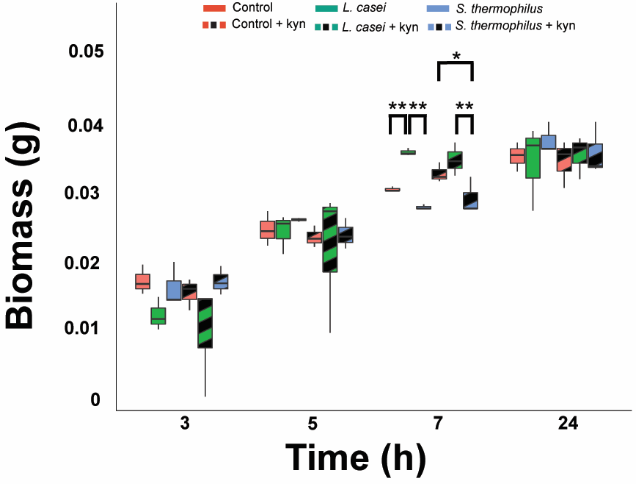

**Supplementary figure 5: Total biomass comparison of the communities**

The boxplots indicate the total biomass produced at each sampling point. Significance as calculated by controlling the false discover rate according the Benjamini-Hochberg procedure is indicated with asterisk; *: q < 0.05 and **: q < 0.01.

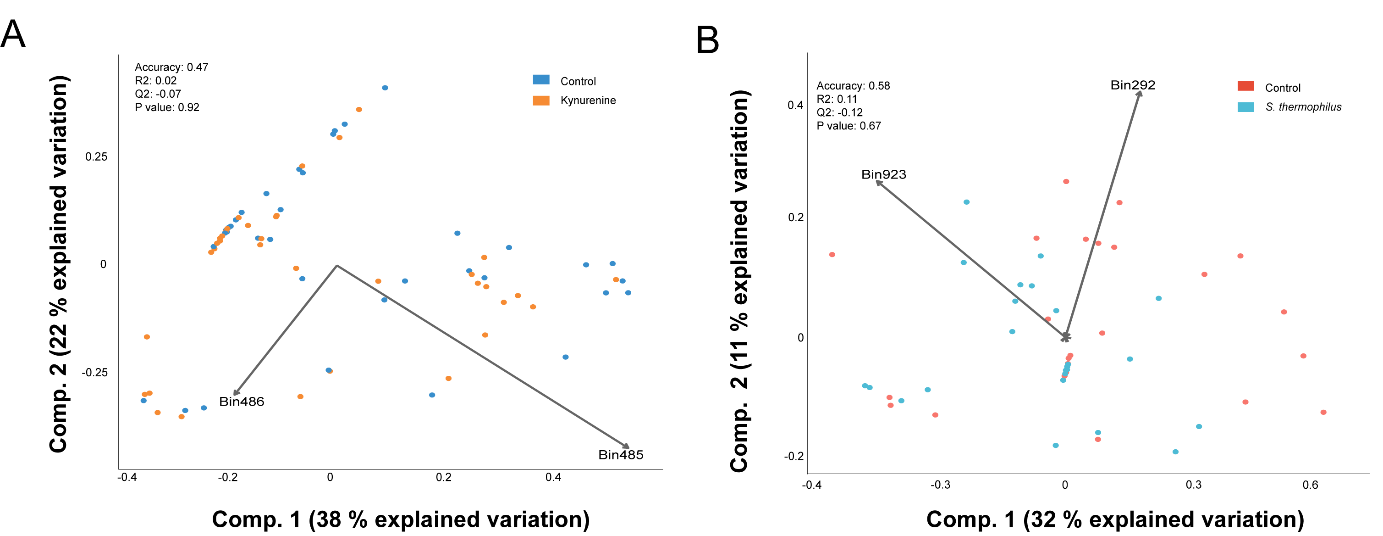

**Supplementary figure 6: partial least square – discriminant analysis of the ^1^H-NMR spectra does not separate the spectra based on kynurenine or *Streptococcus thermophilus* supplementation**

A) Partial least square – discriminant analyses performed with the binned spectra obtained with ^1^H-NMR of the control and kynurenine supplemented communities as input. The bins contributing the most to the separation are indicated with arrows and belong to acetate (bin 485 and 486). B) Partial least square – discriminant analyses performed with the binned spectra obtained with ^1^H-NMR of the control and *Streptococcus thermophilus* supplemented communities as input. The bins contributing the most to the separation are indicated with arrows and belong to an unidentified compound (bin 292) and lactate (bin 923). The accuracy, R2 and Q2 values are statistical parameters which estimate the predictive ability of the model and are calculated via cross validation. The *P*-value is the results of a permutation test performed with the separation distance statistic and 1000 permutations. All parameters are calculated using MetaboAnalyst 5.0.

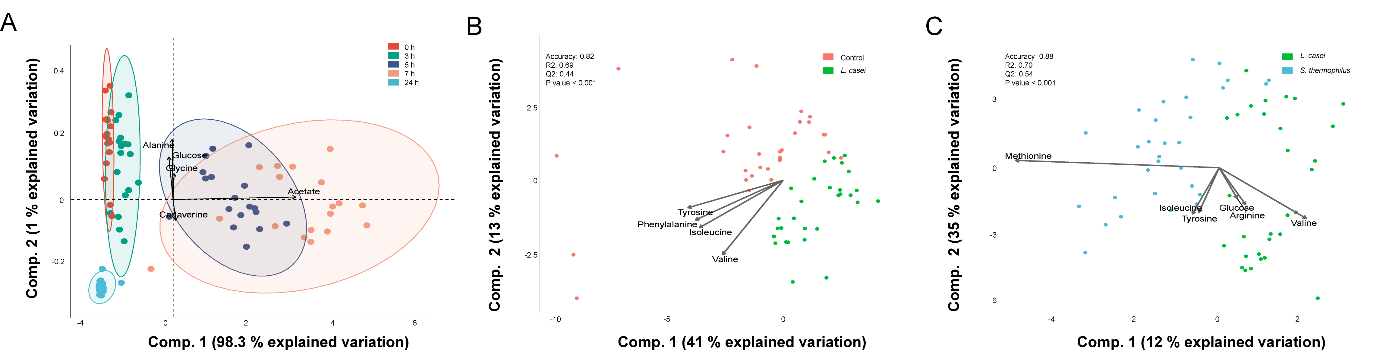

**Supplementary figure 7: ordination analysis of the area under the curve shows a similar separation compared to the binned ^1^H-NMR spectra**

A) Principal component analysis performed with the area under the curve of a representative peak belonging to an identified metabolite as input. The ellipses represent the 95 % confidence interval. The metabolites contributing the most to the separation are indicated with arrows B) Partial least square – discriminant analyses performed with the area under the curve of a representative peak belonging to an identified metabolite of the control and *Lactobacillus casei* supplemented communities as input. The metabolites contributing the most to the separation are indicated with arrows. C) Partial least square – discriminant analyses performed with the area under the curve of a representative peak belonging to an identified metabolite of the *Streptococcus thermophilus* and *Lactobacillus casei* supplemented communities as input. The metabolites contributing the most to the separation are indicated with arrows. For panel A and B: the accuracy, R2 and Q2 values are statistical parameters which estimate the predictive ability of the model and are calculated via cross validation. The *P*-value is the results of a permutation test performed with the separation distance statistic and 1000 permutations. All parameters are calculated using MetaboAnalyst 5.0.

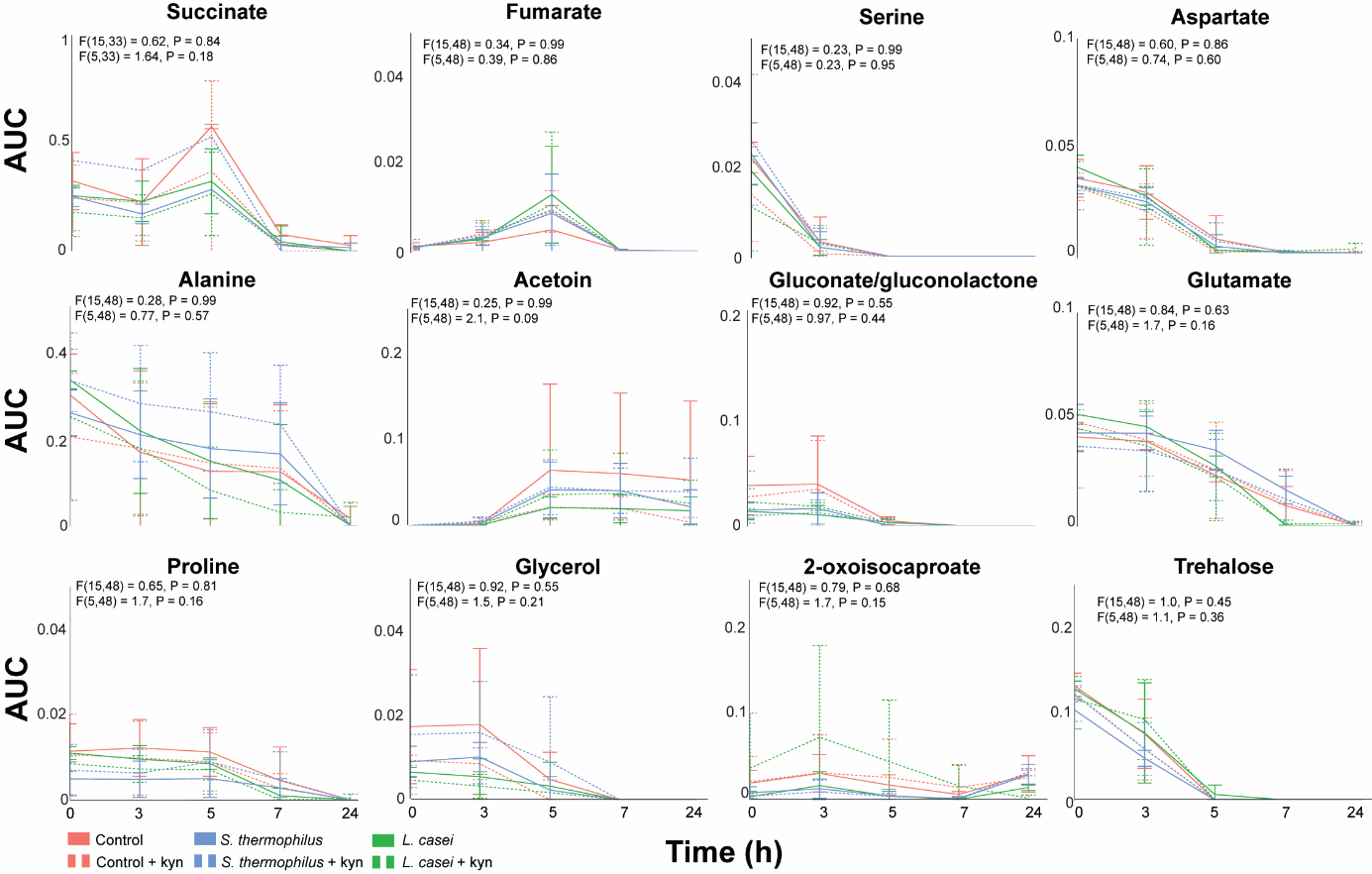

**Supplementary figure 8: similar dynamic metabolic profiles between the kynurenine and probiotic supplemented communities**

Non significantly different dynamic metabolic profiles. The error bars indicate the standard deviation per timepoint. Differences are investigated via repeated measure two way ANOVA. The F statistic and P value for the factor condition and for the interaction between time and condition are indicated for each metabolite.
